# *BRCA* mutational status shapes the stromal microenvironment of pancreatic cancer linking CLU^+^ CAF expression with HSF1 signaling

**DOI:** 10.1101/2021.08.18.456576

**Authors:** Lee Shaashua, Meirav Pevsner-Fischer, Gil Friedman, Oshrat Levi-Galibov, Subhiksha Nandakumar, Reinat Nevo, Lauren E. Brown, Wenhan Zhang, Yaniv Stein, Han Sang Kim, Linda Bojmar, William R. Jarnagin, Nicolas Lecomte, Roni Stok, Hend Bishara, Rawand Hamodi, Ephrat Levy-Lahad, Talia Golan, John A. Porco, Christine A. Iacobuzio-Donahue, Nikolaus Schultz, David Lyden, David A. Tuveson, David Kelsen, Ruth Scherz-Shouval

## Abstract

Cancer-associated fibroblasts (CAFs) give rise to desmoplastic stroma, which supports tumor progression and metastasis, and comprises up to 90% of the tumor mass in pancreatic cancer. Recent work by us and others has shown that CAFs are transcriptionally rewired by adjacent cancer cells to form heterogeneous subtypes. Whether this rewiring is differentially affected by different driver mutations in cancer cells is largely unknown. Here we address this question by dissecting and comparing the stromal landscape of *BRCA*-mutated and *BRCA* Wild-type (WT) pancreatic ductal adenocarcinoma (PDAC). We comprehensively analyze PDAC samples from a cohort of 42 patients by laser-capture microdissection, RNA-sequencing and multiplexed immunofluorescence, revealing different CAF subtype compositions in germline *BRCA*-mutated *vs. BRCA*-WT tumors. In particular, we detect an increase in a subset of Clusterin (CLU)-positive CAFs in *BRCA*-mutated tumors. We further unravel a network of stress responses upregulated in *BRCA*-mutated tumors. Using cancer organoids and cell co-cultures, we show that the transcriptional shift of pancreatic stellate cells into CLU^+^ CAFs is mediated through activation of heat-shock factor 1 (HSF1), the transcriptional regulator of *Clu*. Our findings unravel a new dimension of stromal heterogeneity, influenced by germline mutations in cancer cells, with direct translational implications for clinical research.

**Significance:** *BRCA1*/*2* mutations initiate some of the deadliest cancers, yet the fibroblastic microenvironment of *BRCA*-mutated cancers remains uncharted. Our work addresses a major unsolved question – to what extent is the tumor microenvironment determined by cancer mutations? We find that *BRCA* mutations in the cancer cells affect the composition of CAFs in PDAC. These findings have direct implications for diagnosis and for efforts to exploit CAFs for therapy.

## Introduction

Pancreatic ductal adenocarcinoma (PDAC) is one of the most aggressive cancer types, with a 10% 5-year survival rate [1]. Major contributors to this aggressiveness are cancer-associated fibroblasts (CAFs) [2, 3]. CAFs comprise up to 90% of the cellular tumor microenvironment (TME) in PDAC, and promote tumorigenesis by elevating proliferation, invasion, and chemoresistance of cancer cells, and by remodeling the extracellular matrix (ECM) [4-6]. CAFs are functionally and phenotypically heterogeneous, and are composed of multiple subpopulations [7-10]. In PDAC, CAFs were divided into three subtypes: a myofibroblastic subtype that expresses α-smooth-muscle-actin (αSMA; termed myCAF), an inflammatory subtype that expresses IL-6 and leukemia inhibitory factor (LIF; termed iCAF), and an antigen-presenting subtype that expresses MHC class II (apCAF) [11-13]. Another study described four CAF subtypes with distinct functional features and prognostic impact [9], and a single-cell analysis of human PDAC identified 8 fibroblast clusters [14]. Moreover, cancer-associated mesenchymal stem cells were shown to secrete Granulocyte-macrophage colony-stimulating factor (GM-CSF), acting as CAFs to support PDAC tumor progression [15], while CAFs of pancreatic stellate cells (PSC)-origin were demonstrated to regulate specific ECM features and to contribute to tumor stiffness [16]. Most recently, single-cell analysis of human PDAC identified a subset of LRRC15^+^ CAFs and showed a correlation between elevated levels of this subset and poor response to anti-PD-L1 therapy [17].

CAFs are genomically stable, and rarely have copy number alterations or somatic mutations leading to loss of heterozygosity [18]. Yet, they are transcriptionally heterogeneous [8, 12]. This heterogeneity is driven by different external cues received from neighboring cells and local environmental conditions [19, 20]. For example, hypoxia was shown to induce a pro-glycolytic transcriptional program in CAFs [21], and a metabolic switch from oxidative phosphorylation to glycolysis was also shown in response to TGFβ and PDGF in an IDH3α-mediated mechanism [22]. The stress-induced transcriptional regulator Heat Shock Factor 1 (HSF1) was shown, by us and others, to play a key role in shaping CAF transcription in diverse human carcinomas, including breast, lung, gastric, and colon cancer [23-27]. HSF1 orchestrates a transcriptional program in fibroblasts that enables their reprogramming into CAFs and promotes malignancy by TGFβ and SDF1, YAP/TAZ signaling, and exosome-mediated secretion of THBS2 and INHBA [23-27]. CAF heterogeneity was also proposed to stem from different cells of origin giving rise to CAFs, including tissue resident fibroblasts, mesenchymal stromal cells, pericytes, and adipocytes [8, 15, 28-32]. For example, bone-marrow derived CAFs in breast cancer were shown to express high levels of *Clusterin* (*Clu*), and exhibit a distinct transcriptional profile compared to tissue resident CAFs [31]. However, it is not known whether different germline mutations in the cancer cells lead to differential rewiring of CAFs and contribute to CAF heterogeneity. Moreover, the extent to which CAF rewiring is driven by genomic alterations in the cancer cells versus environmental stresses is not clear.

In PDAC, a subset of up to 7% of the general population, and up to 15% in certain subgroups (such as patients of Ashkenazi Jewish descent), have germline mutations in the breast cancer-1 (*BRCA1*) and *BRCA2* genes [33, 34], which are part of the DNA damage homologous repair mechanism. *BRCA* mutations are the most prominent germline mutations associated with increased risk of developing pancreatic cancer [35]. Patients carrying these mutations, both in PDAC and in other *BRCA*-associated cancers (e.g. breast cancer), exhibit a higher response rate to platinum-based chemotherapy regimens and PARP-inhibitors, resulting in longer than expected overall survival [34, 36]. Several cell-autonomous mechanisms by which PARP inhibitors affect *BRCA*-mutant (BRCA-mut) cancer cells were suggested [37-39], however, additional non-cell-autonomous factors mediating the efficacy of these treatments may be pivotal in PDAC. Recent studies described distinct immune-microenvironments in *BRCA*-mut cancers, characterized by increased infiltration of T cells and macrophages [40-43]. Since the stromal cells are genomically stable [18], it is suggested that the rewiring of the TME is orchestrated by *BRCA* mutations in the cancer cells. Yet the transcriptional landscape of the fibroblastic microenvironment of *BRCA*-mut PDAC remains uncharted. In breast cancer, we have recently identified two major CAF subtypes expressing either the marker S100A4 (also known as FSP1) or Podoplanin (PDPN) [44]. The ratio of these two CAF subtypes was correlated with *BRCA1/2* mutational status and with disease outcome in *BRCA*-mut breast cancer patients.

Given that CAFs are reprogrammed by the adjacent cancer cells, we hypothesize that different driver mutations will yield different stromal landscapes. Here, we set out to test this hypothesis in a comprehensive cohort of 42 *BRCA*-mut and *BRCA*-WT pancreatic cancer patients. Using three CAF markers – Clusterin (CLU), αSMA, and MHC class II – we identify three mutually exclusive CAF subtypes in primary pancreatic tumor resection specimens, and show that the ratio between these CAF subtypes is altered in *BRCA-*mut tumors compared to *BRCA*-WT tumors. We apply laser capture microdissection (LCM) followed by RNA-sequencing to define stromal transcriptional signatures unique to *BRCA*-mut *vs. BRCA*-WT tumors. We characterize these signatures by multiplexed immunofluorescence (MxIF) and find distinct stress response activation patterns in *BRCA*-mut *vs. BRCA*-WT tumors. In particular, we find that HSF1 is upregulated in *BRCA*-mut tumors. Using cancer organoids and cell co-cultures, we show that HSF1 transcriptionally activates *Clu* and mediates the transcriptional shift of PSCs into CLU^+^ CAFs. Our findings portray distinct stromal compositions in *BRCA*-mut and *BRCA*-WT PDAC tumors that may serve as targets for early detection and for PDAC therapy.

## Results

### *BRCA*-mut and *BRCA*-WT tumors exhibit distinct CAF compositions

To dissect the stroma of *BRCA*-mut PDAC in comparison with that of *BRCA*-WT PDAC, we assembled a clinical cohort of 42 patients (27 *BRCA*-WT and 15 germline *BRCA*-mut; see Supplementary Table 1). Formalin-fixed primary tumor resection tissue, and deeply annotated demographic, clinical and pathologic data were collected for all patients in the study. In addition, genomic (MSK-IMPACT™) data and fresh-frozen tumor tissue was collected for a subset of patients. PDAC CAFs are comprised of distinct subtypes, marked by the expression of distinct proteins [9, 11, 12]. To test whether CAF compositions are affected by the germline mutational status of the cancer cells we assessed the distribution of several CAF markers in primary tumor resections from *BRCA*-WT and *BRCA*-mut PDAC patients. We found three markers that marked discrete CAF subtypes – α-smooth-muscle-actin (αSMA), clusterin (CLU) and human leukocyte antigen DR isotype (HLA-DR) (an MHC class II molecule; Figure 1A-B and Figure S1A-C). αSMA is a well-known myofibroblastic marker in various carcinomas, including PDAC [45]. *Clu* was shown to be expressed by αSMA^low^ CAFs in breast and pancreatic cancer [8, 12, 31]. MHC-II was recently suggested as a marker of apCAFs in both breast and pancreatic cancers [8, 11]. Immunohistochemical analysis (IHC; Figure 1A) and multiplexed immunofluorescence (MxIF; Figure 1B) staining demonstrated a segregation of αSMA^+^, CLU^+^ and HLA-DR^+^ CAFs in PDAC. Automated image analysis quantifying the relative abundance of these proteins in stromal cells (CD45^-^Cytokeratin^-^) in a subcohort of 7 *BRCA-*mut and 6 *BRCA-*WT tumors confirmed that the three CAF markers clearly mark discrete CAF subtypes (Figure S1A-C).

**Figure 1.**
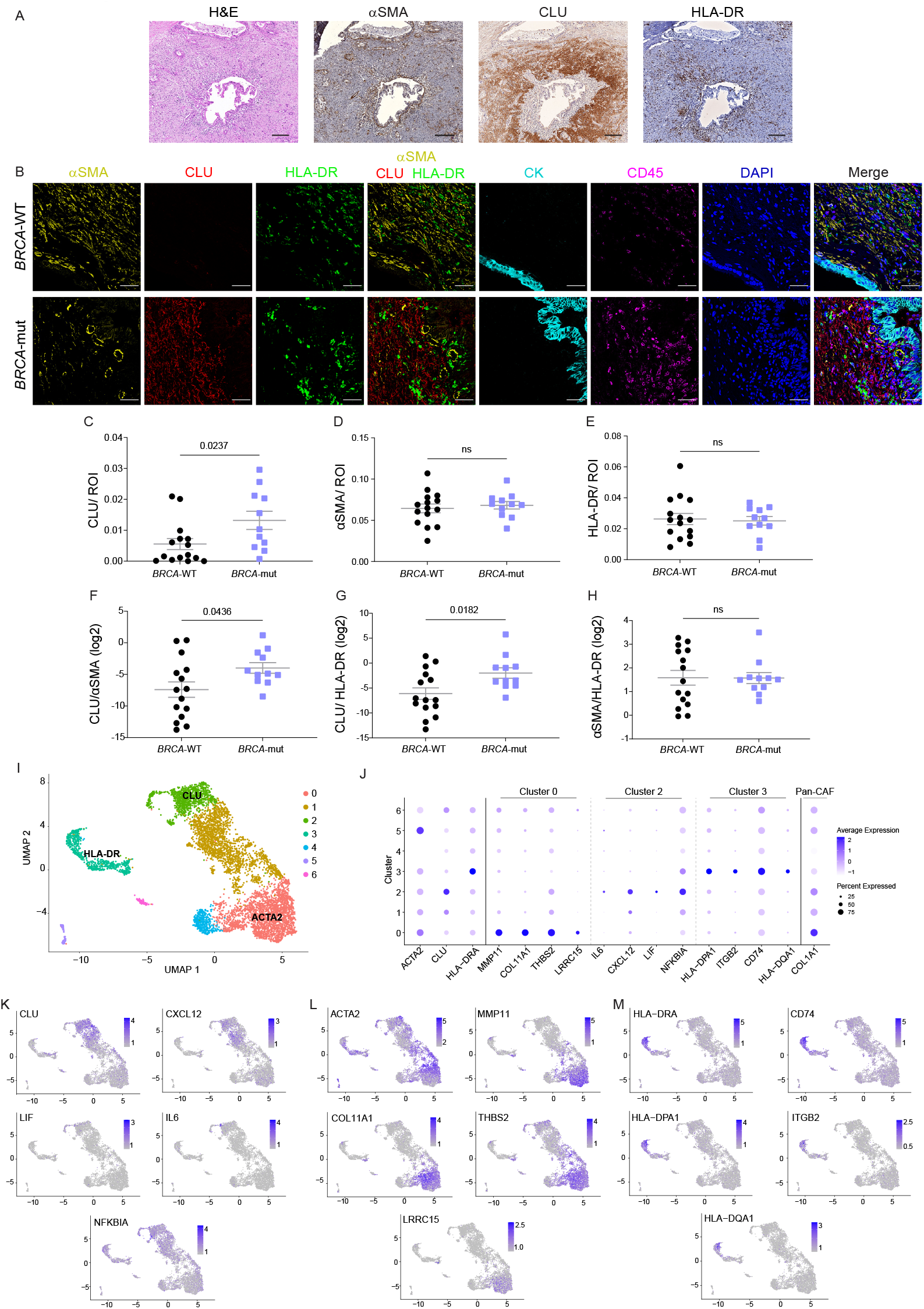
CAF compositions change between *BRCA*-WT and *BRCA*-mut PDAC tumors. Formalin-fixed Paraffin-embedded (FFPE) tumor sections from *BRCA*-mut and *BRCA*-WT PDAC patients were stained for H&E, IHC, and MxIF. (A) IHC was performed for αSMA, CLU and HLA-DR (Scale bar, 200μm). Representative images are shown. (B-H) MxIF was performed using antibodies for the depicted proteins. DAPI was used to stain nuclei. Scale bar, 50μm. Representative images are shown in (B). Images were analyzed using ImageJ software, CD45^-^ CK^-^ regions were defined as regions of interest (ROIs) and the area stained by each CAF marker was calculated, divided by the ROI and averaged for each patient sample (C-E). Mann Whitney test was performed. The ratio of the different CAF subtypes is shown in (F-H) and was analyzed using Student’s t-test. Data are presented as Mean ± SEM. ns marks p-values greater than 0.05. For IHC and H&E staining n=2, and for MxIF staining n=11 *BRCA*-mut and n=15 *BRCA*-WT. (I-M) A single-cell human PDAC dataset [14] was reanalyzed using the Seurat R toolkit. (I) Uniform Manifold Approximation and Projection (UMAP) map of 6,405 cells, color-coded for the indicated cell clusters defined by a local moving clustering algorithm. The clusters that differentially express *ACTA2, CLU* and *HLA-DR* are indicated. (J) Dot plot visualization of gene expression of the indicated CAF markers. (K-M) Single-cell expression level of CAF markers on the UMAP shown in (I). Marker genes of *CLU* (K), *ACTA2* (L) and *HLA-DR* (M) clusters are represented.

Next, we asked whether the global composition of immune cells, cancer cells and CAFs is different between *BRCA-*mut and *BRCA-*WT tumors. We quantified the number of CD45^+^ (immune) cells by MxIF and found no differences between the different genotypes (Figure S1D). Then, we used an artificial intelligence image analysis algorithm to classify different cell populations in hematoxylin & eosin–stained FFPE sections from patients. We found no significant differences in the percentage of CAF-rich, cancer-rich, and immune-rich regions between *BRCA-*mut and *BRCA-*WT tumors (Figure S1E-H). We then evaluated each subpopulation of CAFs separately, by quantifying CLU^+^, MHC-II^+^, and αSMA^+^ CAF staining in a subcohort of 26 human PDAC patients including 15 *BRCA-*WT and 11 *BRCA-*mut patients. While αSMA^+^ CAFs and HLA-DR^+^ CAFs did not differ between the genotypes, CLU^+^ CAFs were significantly more abundant in *BRCA*-mut tumors (Figure 1C-E). Moreover, the ratio between CAF subtypes was different between *BRCA-*mut and *BRCA-*WT tumors (Figure 1F-H). Specifically, the ratio of CLU^+^/αSMA^+^ CAFs and the ratio of CLU^+^/HLA-DR^+^ CAFs was higher in *BRCA*-mut tumors compared to *BRCA*-WT tumors (Figure 1F-G), suggesting that germline mutations in the cancer cells alter tumor CAF compositions.

To examine whether this characteristic is different between *BRCA1* and *BRCA2* mutant tumors, we compared the relative abundance of CLU^+^ CAFs and the CLU^+^/αSMA^+^ ratio in *BRCA1 vs. BRCA2* mutant patients. We found no significant differences (Figure S1I-J), implying that the changes in CAF compositions between *BRCA*-mut and *BRCA*-WT tumors are shared between different *BRCA* mutations.

Several recent studies associated CLU expression with neoadjuvant therapy. One study reported elevated expression of CLU following neoadjuvant therapy [46], and another suggested that stromal expression of CLU is a predictive marker of the response to neoadjuvant therapy [47]. We therefore compared CLU^+^/αSMA^+^ CAF ratios in neoadjuvant-treated *vs*. non-treated patients. We found that the ratio of CLU^+^/αSMA^+^ was not affected by treatment (Figure 1SK). Both treated and non-treated patients had higher CLU^+^/αSMA^+^ CAF ratios in *BRCA*-mut patients compared to WT, supporting the notion that CAF distribution is driven by the tumor genotype and is not altered by neoadjuvant treatment regimens (See Table 1 for clinical information).

Next, we sought to explore whether the CAF subtypes we characterized using protein markers could also be identified at the transcriptional level. To that end we utilized a recently published single-cell RNA-seq dataset of human PDAC from Peng et al. [14], and reanalyzed the data from all tumor samples in this dataset (i.e. excluding cells from normal controls). We used the Seurat R toolkit [48] to analyze all the cells that were defined as fibroblasts or stellate cells in the original dataset, and excluded *MCAM* positive cells (a pericyte marker). Unbiased clustering revealed 7 distinct CAF subtypes (Figure 1I-J, Supplementary Table 2). *ACTA2, CLU*, and *HLA-DR* were significantly differentially expressed in distinct clusters, supporting our MxIF analysis and suggesting that not only at the protein level, but also at the transcriptional level, these genes mark discrete CAF populations. Pathway analysis using the Metascape software [49] of the top differentially expressed genes for each of these 3 clusters revealed distinct transcriptional programs that correspond with the previously described subtypes – myCAF (*ACTA2*), iCAF (*CLU*), and apCAF (*HLA-DR*; Figure 1J-M). The *ACTA2*^*+*^ (αSMA) cluster was enriched for myofibroblastic pathways such as ECM remodeling (Collagens and MMPs), wound healing (*INHBA, THBS2*), smooth muscle contraction (*ACTA2 and TPM* genes), and cell-substrate adhesion (*LRRC15, ITGB5*) [11, 12, 17, 25]. The *CLU*^*+*^ cluster expressed inflammatory genes (*IL-6, CXCL12, CXCL1, NFKBIA*), as well as genes involved in ECM remodeling (*LIF, COL14A1, HAS1*) and angiogenesis regulation (*C3, IL6*) [11-13]. The *HLA-DR* cluster was enriched for antigen presentation (variety of *HLA* genes), and for T cell activation (*ITGB2, S100A8*). These results indicate that CLU is a marker of a discrete CAF subset in PDAC, characterized by an inflammation-associated gene signature.

### *BRCA*-WT and *BRCA*-mut stroma exhibit distinct transcriptional signatures

To directly map the transcriptional landscapes of *BRCA*-WT and *BRCA*-mut stroma, we employed laser capture microdissection (LCM) followed by RNA-sequencing on CAF-rich regions from 12 patients (5 *BRCA*-mut and 7 *BRCA*-WT; Supplementary Table 3). Differential expression analysis revealed 30 upregulated and 10 down-regulated genes in *BRCA*-mut *vs. BRCA*-WT patients (Figure 2A). Pathway analysis showed enrichment of genes involved in ECM remodeling and proteolysis (*MUC5B, SERPINA1, A2ML1, S100A2, GREM1*), wound healing (*TNC, CD177, WFDC1*), muscle contraction (*DES, KCNMA1, OXTR, CEMIP*), and regulation of cell growth (*CRABP2, ROS1, WFDC1*) in *BRCA*-mut *vs. BRCA*-WT patients (Figure 2A and Supplementary Table 4). Genes involved in T-cell activation and migration (*IRF4, TBX21, CXCL9*) and tyrosine kinase signaling (*STAP1, FLT3*) were downregulated in *BRCA*-mut *vs. BRCA*-WT patients (Figure 2A and Supplementary Table 4). To exclude the possibility that the observed differential expression of immune-related genes is due to higher immune-cell contamination of the dissected CAF-rich regions in *BRCA*-WT stroma, we applied CIBERSORT, a computational deconvolution tool that estimates the relative abundance of individual cell types in a mixed cell population based on bulk RNA-seq profiles [50]. The distribution of all tested immune cell populations was similar between *BRCA*-WT and *BRCA*-mut stroma (Figure S2A and Supplementary Table 5), supporting the conclusion that even if immune cells have infiltrated into the dissected stromal regions, the observed differential expression patterns most likely originate from CAFs.

**Figure 2.**
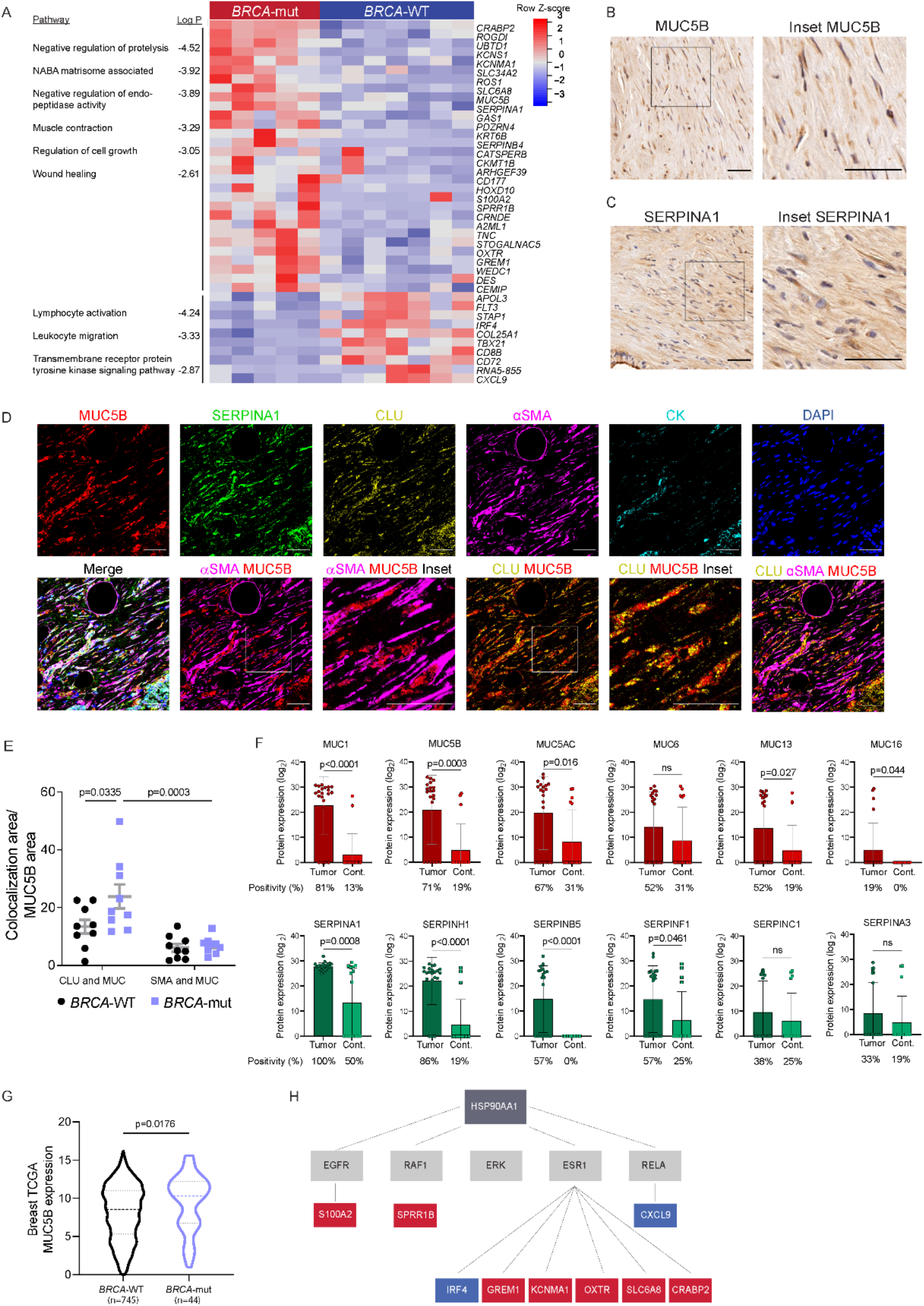
The transcriptional profile of *BRCA*-mut stroma is different than that of *BRCA*-WT stroma. CAF-rich regions of fresh-frozen tumor sections from 7 *BRCA*-WT and 5 *BRCA*-mut PDAC patients were laser-capture-microdissected and analyzed by RNA-seq. (A) Heatmap showing hierarchical clustering of differentially expressed (DE) genes in CAF-rich regions from *BRCA*-mut and *BRCA*-WT samples. Pathway analysis was performed using Metascape; Selected significant pathways (p<0.05) are shown (see full list in Supplementary Table 4). (B-C) FFPE tumor sections from 2 PDAC *BRCA*-mut patients were stained by IHC for MUC5B and SERPINA1. Representative images are shown. Scale bar, 50μm (D-E) FFPE tumor sections from 9 *BRCA*-mut and 9 *BRCA*-WT PDAC patients were stained by MxIF using antibodies for the indicated proteins. DAPI was used to stain nuclei. Scale bar, 50μm. Representative images are shown in (D). MUC5B and SERPINA1 protein levels were quantified by ImageJ software and the area stained by each protein and CAF marker was measured. Quantification of MUC5B colocalization with CLU and αSMA, analyzed by two-way ANOVA and presented as mean ± SEM is presented in (E). (F) Human pancreatic adenocarcinoma tissue-derived exosomal proteomes (n=21) and non-tumor adjacent tissue-derived exosomal proteomes (n=16) were analyzed by liquid chromatography with tandem mass spectrometry (LC-MS/MS). Proteins found in >15% of pancreatic cancer exosomes were compared to pancreatic adjacent tissue-derived exosomes. Log_2_ protein expression of the indicated proteins is presented. P values were calculated by Welch’s t-test for the comparison of expression level and Fisher’s exact test for the comparison of positivity. Data are expressed as mean ± SEM. (G) Expression of *MUC5B* in *BRCA*-mut (n=44) and *BRCA*-WT (n=745) breast cancer patients from the TCGA. Data were analyzed by Student’s t-test and are presented by violin plots. (H) DE genes were analyzed by Ingenuity software using the causal network tool. Schematic representation of the predicted network is presented. Upregulated and downregulated genes in *BRCA*-mut patients are marked in red and blue, respectively; predicted regulators are marked in grey.

We next set to validate some of the differentially expressed genes at the protein level. We chose to focus on *MUC5B* and *SERPINA1*, two of the most significantly upregulated genes in *BRCA*-mut CAFs. These genes encode secreted proteins, which were previously proposed to serve as prognostic biomarkers of pancreatic neoplasms based on proteomic analysis of pancreatic fluids. SERPINA1 levels were elevated in PanIN3 lesions [51], and MUC5B was identified in pancreatic main duct fluid collected at the time of surgical resection [52]. IHC staining of tumor sections from PDAC patients confirmed that MUC5B and SERPINA1 are expressed in PDAC stroma (Figure 2B-C). MxIF staining showed similar protein levels of SERPINA1 and MUC5B in *BRCA*-mut *vs. BRCA*-WT tumors, when analyzing all stromal cells. However, further analysis of MUC5B expression within the different CAF subtypes revealed that MUC5B is expressed in CLU^+^ CAFs and significantly less so in SMA^+^ CAFs, and that the localization of MUC5B within CLU^+^ CAFs is significantly higher in *BRCA*-mut patients (Figure 2D-E). To test whether these proteins are indeed secreted by PDAC human tumors, we assessed the exosomal content of 21 PDAC specimens and 16 normal adjacent controls in an independent patient cohort (see Methods and Supplementary Table 6). We detected multiple mucin and serpin proteins that were highly expressed in tumor exosomes compared to normal adjacent tissue-derived exosomes (Figure 2F and Supplementary Table 6). Specifically, MUC5B was detectable in 71% of PDAC-derived exosomes, compared to 19% of adjacent pancreatic tissue-derived exosomes. SERPINA1 was evident in 100% of PDAC-derived exosomes, but was less specific as it was found in 50% of the control tissues (Figure 2F and Supplementary Table 6). We next set out to validate the higher stromal expression of MUC5B in *BRCA*-mut tumors in an additional dataset. Since no published PDAC datasets have a sufficient number of *BRCA*-mut patients, we turned to The Cancer Genome Atlas (TCGA) breast cancer dataset, and analyzed the expression of *MUC5B* in *BRCA*-mut *vs. BRCA*-WT tumors. *MUC5B* expression was significantly higher in *BRCA*-mut breast cancer patients compared to *BRCA*-WT (Figure 2G), suggesting that this protein may serve as a Pan-*BRCA* stromal marker.

To identify potential upstream regulators of the *BRCA*-associated CAF transcriptional program we analyzed our differentially expressed gene dataset using the Causal Network tool in the Ingenuity Pathway Analysis (IPA) software [53]. This analysis highlighted heat shock protein 90α gene, *HSP90AA1*, as a potential upstream regulator of multiple genes in our network (Figure 2H). HSP90α is a stress-induced chaperone. Previous studies have reported a role for HSP90 in PDAC progression [54], and synergistic effects of CLU and HSP90α in promoting epithelial-to-mesenchymal transition and metastasis in breast cancer [55]. As both *CLU* and *HSP90AA1* are regulated by HSF1 [56, 57], the master transcriptional regulator of the heat shock response, we hypothesized that HSF1 may be orchestrating these *BRCA*-mut-induced transcriptional changes in the stroma.

### Activation of stromal HSF-1 is elevated in *BRCA*-mut PDAC tumors

Work by us and others has shown indispensable roles for HSF1 in transcriptional rewiring of fibroblasts into CAFs in various cancer types [23-27]. To test whether HSF1 is differentially activated in *BRCA*-mut *vs. BRCA*-WT CAFs, we performed MxIF staining. HSF1 translocates to the nucleus upon activation, and thus its nuclear localization serves as a proxy for its activation (Figure 3A-C). Comparing 16 *BRCA*-mut tumors with 20 *BRCA*-WT tumors from our patient cohort, we found significantly higher activation of HSF1 in *BRCA*-mut stroma compared to *BRCA*-WT stroma (Figure 3C). We were next curious to see whether other stress responses are also activated in *BRCA*-mut stroma, possibly due to DNA-damage-induced stress, or whether this phenomenon was specific to HSF1. To portray the stress network in PDAC, we stained for five additional stress-induced transcription factors (TFs): X-box binding protein 1 (XBP1) [58] and Activating Transcription Factor 6 (ATF6 [59]; ER-stress response); Hypoxia-inducible factor 1-α (HIF1α ; Hypoxia) [21], Nuclear factor erythroid-2-related factor 2 (NRF2; oxidative stress) [60], and Activating Transcription Factor 4 (ATF4 [61]; the integrated stress response; Figure 3A-E). While none of these additional stress-activated TFs showed significant differential activation (Figure S3A-E), a significant crosstalk between all these stress pathways was evident. All pairs of stress-TFs exhibited higher co-activation (per-patient) in *BRCA*-mut tumors compared to *BRCA*-WT tumors (Figure 3D-E), suggesting that a network of stress responses is activated in the stroma of *BRCA*-mutated PDAC.

**Figure 3.**
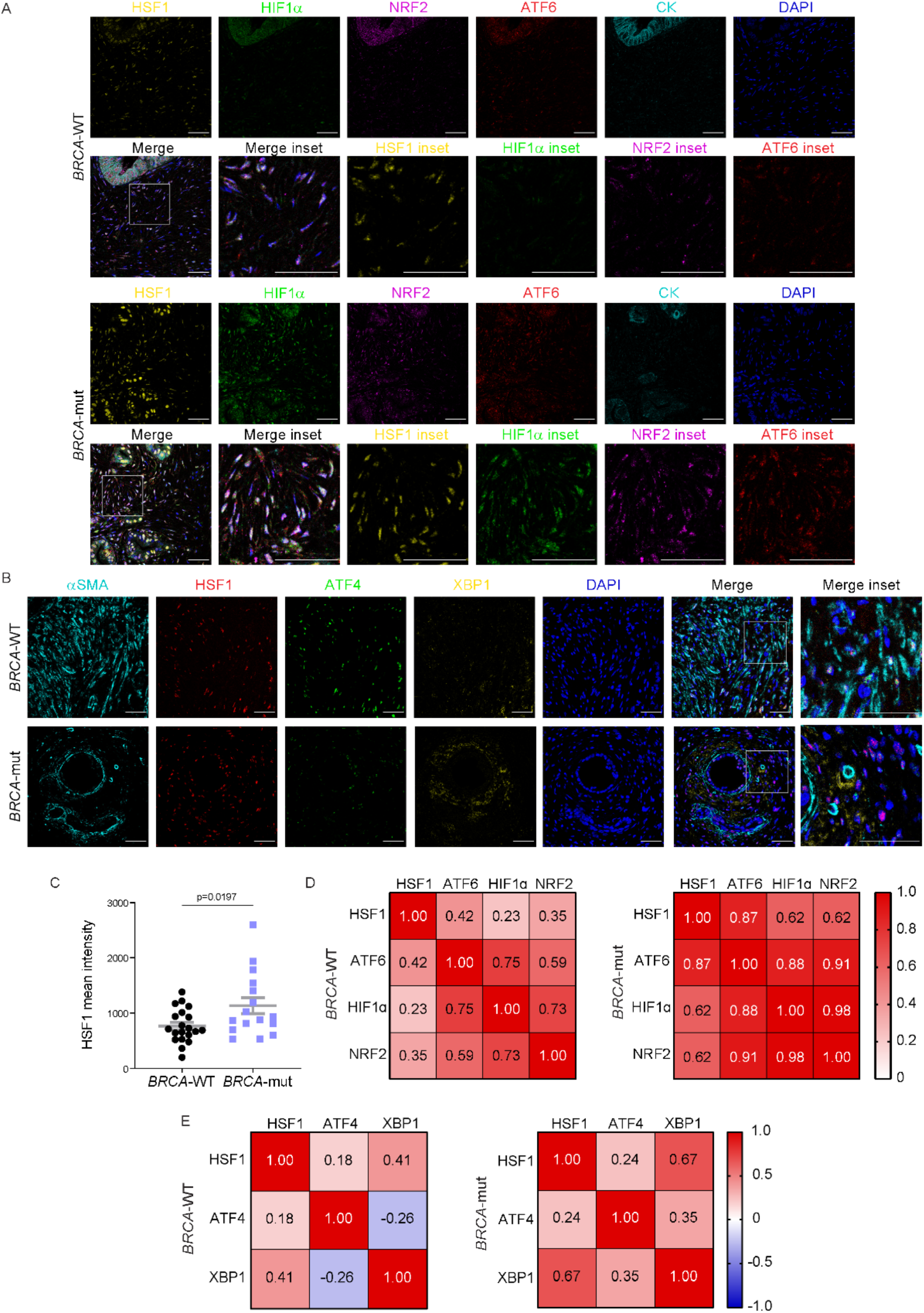
A network of stress responses is activated in *BRCA*-mut stroma. FFPE tumor sections from *BRCA*-mut and *BRCA*-WT PDAC patients were stained by MxIF using antibodies for the indicated proteins. (A-B) Representative images are shown. DAPI was used to stain nuclei. Scale bar, 50μm. (C) Quantification of HSF1 mean intensity within all stromal cells in *BRCA*-mut (n=16) and *BRCA*-WT samples (n=20). 3-5 images per patient were analyzed using ImageJ software, HSF1 staining intensity was averaged within patients, and is presented as mean (across patients) ± SEM. (D-E) Pearson correlation matrices of stress TF coactivation in *BRCA*-mut (n=8 for D; n=13 for E) *vs. BRCA*-WT (n=11 for D; n=17 for E) patients.

### HSF1 upregulates CLU/αSMA ratio in *BRCA*-mut tumors

CLU is an extracellular chaperone transcriptionally regulated by HSF1, and upregulated in response to DNA damage [62, 63]. CLU was shown to play a critical role in promoting pancreas regeneration and tumorigenesis [64, 65]. Supported by our findings of higher HSF1 activation and CLU/αSMA ratios in *BRCA*-mut tumors, we hypothesized that HSF1 may affect *BRCA*-associated CAF compositions through transcriptional regulation of stromal gene expression and specifically regulation of *CLU* expression. To test this hypothesis, we first assessed the correlation between HSF1 activation and CLU/αSMA ratio in our clinical cohort. We found that HSF1 activation is correlated with CLU/αSMA ratio only in *BRCA*-mut patients and not in *BRCA*-WT patients (Figure 4A-B). Next we asked whether *CLU* expression is HSF1-dependent. To this end, we measured mRNA expression of *Clu* in primary PSCs isolated from WT and *Hsf1* null mice (Figure 4C). We found that the expression of *Clu* was significantly lower in *Hsf1* null PSCs compared to WT PSCs, while the expression of other CAF markers, such as *Acta2* and *Il6*, was not altered (Figure 4E-F). *Muc5B* showed somewhat reduced expression but this result was not significant (Figure 4D).

**Figure 4.**
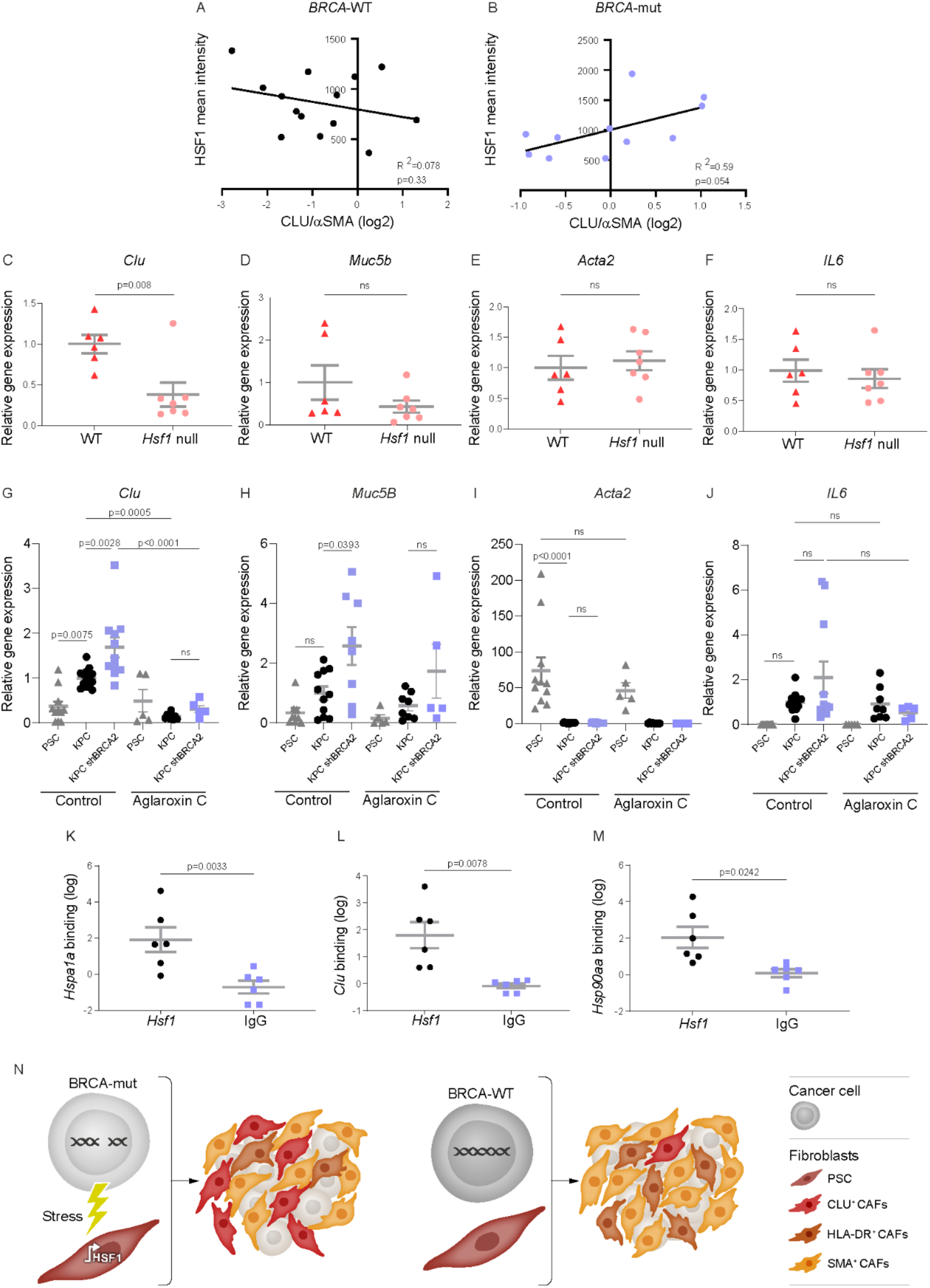
HSF1 directly regulates *Clu* expression in BRCA-mut tumors. (A-B) FFPE tumor sections from 14 *BRCA*-WT and 11 *BRCA*-mut PDAC patients were stained by MxIF for HSF1, CLU and αSMA. Images were analyzed by ImageJ. Pearson correlation between HSF1 mean intensity and CLU/αSMA ratio in (A) *BRCA*-WT patients, and (B) *BRCA*-mut patients was calculated. (C-F) Expression levels of *Clusterin* (C), *Muc5b* (D) *Acta2* (E) and *Il6* (F) in primary PSCs freshly isolated from WT and *Hsf1*-null mice. (G-I) Immortalized PSCs were seeded in Matrigel for 4 days in 10% FBS in DMEM. Conditioned media (CM) from PSCs or from KPC cells in which *Brca2* was silenced by shRNA or non-targeting shControl (KPC) was then added for an additional 4 days. 3nM of the HSF1 inhibitor, CMLD011866 (Aglaroxin C), or PBS control was added to the conditioned media (CM) every 2 days, for 8 days, after which the expression levels of *Clusterin* (G), *Muc5b* (H) *Acta2* (I), and IL6 were measured by qRT-PCR. (K-M) Immortalized PSCs were cultured with or without shControl KPC CM for 24 hours. ChIP-PCR was performed for putative heat-shock elements of *Hsp1a1* (K), *Clu* (L), and *Hsp90aa* (M) following pulldown with anti-HSF1 antibody compared to IgG control. Data are presented as mean ± SEM in C-M. (N) Schematic representation of the proposed model. *BRCA* mutations in the cancer cells provoke a stressful microenvironment. *HSF1* activation in the adjacent PSCs leads to their rewiring into CLU^+^ CAFs.

To characterize the effect of *BRCA* mutations on HSF1-dependent *Clu* upregulation, we employed shRNA for *Brca2* in KPC cells (mimicking BRCA2 loss-of-function) or non-targeting (NT) control (Figure S4). We chose to target *Brca2* rather than *Brca1* since mutations in *BRCA2* are more prevalent than in *BRCA1* in PDAC, and were found in 73% of our *BRCA*-mut cohort. Immortalized PSCs were cultured in 3D matrigel domes for four days in growth medium and four additional days in the presence of conditioned media (CM) from KPC pancreatic cancer cell-organoids transduced with shRNA for *Brca2* or NT control. CM from the untreated PSCs served as control for both conditions (Figure 4G-J). These growth conditions were previously shown to suppress the myCAF phenotype [12] and indeed we found that *Acta2* expression was abolished by the addition of KPC CM (Figure 4 I). In stark contrast, the expression of *Clu* and *Muc5b* was induced by addition of KPC CM, and silencing of *Brca2* in the KPCs led to a further, significant induction of both *Clu* and *Muc5b* expression (as compared to CM from NT controls or from PSCs; Figure 4G-H). *Il6* expression was also induced by KPC CM, though this induction was not statistically significant (Figure 4J). To test whether the cancer-induced upregulation of *Clu* and *Muc5b* is HSF1-dependent, we added to these cultures the synthetic small molecule CMLD011866 ((-)-aglaroxin C) [66-68]. This compound is a pyrimidinone variant of the rocaglate/flavagline natural product class, recently shown by us to inhibit HSF1 activity [24, 68]. Aglaroxin C was added to the CM every two days and the expression of *Clu, Muc5B, Acta2 and Il6* was measured (Figure 4G-J). Treatment with Aglaroxin C abolished the induction of *Clu* and *Muc5b* expression, suggesting that this induction is HSF1-dependent, and that HSF1 regulates the expression of these genes. Finally, to examine if *Clu* and *Hsp90aa* are direct target genes of HSF1 in our system, we exposed PSCs to KPC-CM and performed chromatin immunoprecipitation (CHIP) with anti-HSF1 antibodies followed by qPCR with primers for heat-shock elements on the DNA of *Clu and Hsp90aa. Hspa1a*, a well-known HSF1-target gene, served as control (Figure 4K-M). Both *Clu* and *Hsp90aa* were significantly enriched in the HSF1-bound fraction compared to IgG control, demonstrating direct regulation of these genes by HSF1 in cancer-conditioned PSCs (Figure 4K-M). Overall, our findings suggest that *BRCA*-mut cancer cells promote a stressful TME that leads to the activation of HSF1 in PSCs. These PSCs are reprogrammed into CLU^+^ CAFs, resulting in a different stromal landscape in *BRCA*-mut compared to *BRCA*-WT PDAC tumors (Figure 4N).

## Discussion

Accumulating evidence over the past few years unraveled vast heterogeneity of CAFs in the TME [7, 8, 69, 70]. This heterogeneity was proposed to stem from different cells of origin giving rise to CAFs [28-32, 71], and from transcriptional rewiring driven by different external cues received from neighboring cells and local environmental conditions [19, 20]. The contribution of germline mutations in the cancer cells to stromal rewiring is largely uncharted. Here we find that the stromal microenvironment of *BRCA*-mut PDAC is distinct from that of *BRCA*-WT PDAC. Specifically, we show that human *BRCA*-mut tumors express higher levels of CLU^+^ CAFs, and, consequently, higher CLU^+^/αSMA^+^ and CLU^+^/HLA-DR^+^ CAF ratios. We portray the transcriptional landscapes and potential upstream regulators of CAFs in *BRCA*-mut and *BRCA*-WT tumors from patients, and reveal a network of stress responses activated in *BRCA*-mut-associated stroma. Within this network, we find a specific role for HSF1 as the transcriptional regulator of *Clu*, mediating the transition of PSCs to CLU^+^ CAFs (Figure 4N).

Recent studies by us and others have utilized single-cell RNA-sequencing and imaging technologies to classify CAFs into functional subtypes based on differential expression of cell surface markers and genes. Here we find three CAF subtypes, distinctively marked by αSMA, CLU and HLA-DR (i.e. MHC-II). αSMA is a classic marker for myofibroblasts. MHC-II was only recently discovered to mark a subtype of CAFs [8, 11], referred to as antigen-presenting CAFs, though their actual antigen-presenting activities remain to be elucidated. CLU marks SMA^low^ CAFs in mouse models of breast and pancreatic cancer [8, 12, 31]. Our MxIF analysis showing that CLU marks a discrete population than HLA-DR^+^ CAFs, together with our single-cell transcriptional analysis highlighting inflammatory pathways in *CLU*^*+*^ CAFs, suggest that in human PDAC CLU marks inflammatory CAFs. These CLU^+^ CAFs are significantly upregulated in *BRCA*-mut PDAC compared to *BRCA*-WT. *BRCA*-mut PDAC immune-microenvironments are characterized by increased infiltration of T cells [40-43]. CLU^+^ CAFs may act as mediators of this immune response, serving to recruit T cells and macrophages. Moreover, the concomitant relative reduction in SMA^+^ CAFs most likely contributes to the observed transcriptional changes in ECM-related genes, which in turn can further affect the composition of the immune-microenvironment.

*Clu* is transcriptionally regulated by the stress-induced master regulator, HSF1 [56]. HSF1 has been shown by us and others to play key roles in transcriptional rewiring of fibroblasts into CAFs in various cancers [23-27]. Here we describe for the first-time preferential activation in a specific subtype of cancer, *BRCA*-mut PDAC, and expose a new facet of HSF1’s stromal activities, affecting CAF composition. Preferentially activated in the stroma of *BRCA*-mut PDAC, HSF1 activates *Clu* and leads to induction of CLU^+^ CAFs.

CLU is a molecular chaperone, harboring two isoforms – a nuclear isoform (nCLU) and a secreted one (sCLU). These isoforms were shown to have opposing activities; nCLU is a pro-apoptotic factor, while sCLU is a stress-induced, pro-survival factor. In epithelial cells, sCLU is upregulated by DNA damage [72], it is overexpressed in various cancers [73], and the shift of nCLU to sCLU expression was shown to be associated with progression towards high-grade and metastatic carcinoma in different cancers [74-77]. The nuclear-to-secreted transition of CLU has not been extensively characterized in fibroblasts. In CAFs, we detect the secreted form of CLU, and find this form significantly upregulated in CAFs of *BRCA*-mut PDAC. Previous studies have implicated TGFβ in promoting the transition of CAFs from inflammatory to myofibroblast-like [13]. Moreover, TGFβ was shown to negatively regulate the expression of sCLU in fibroblasts during fibrosis [78, 79]. In line with this, our RNA-seq analysis shows upregulation of the TGFβ inhibitor, Gremlin1 (*GREM1*), in *BRCA*-mut PDAC. Taken together these findings further support the notion that *BRCA* mutations in the cancer cells lead to a stromal shift from TGFβ induced SMA^+^ myofibroblasts to HSF1-induced CLU^+^ fibroblasts.

This work identifies a unique stress-response-network that is activated in *BRCA*-mut stroma. Tumors are stressful environments and stress responses are well known to play important roles in supporting survival of cancer cells. For example, activation of *Nrf2* in cancer cells leads to elevated mRNA translation and mitogenic signaling [60], the endoplasmic-reticulum (ER) stress response was shown to mediate chemoresistance in PDAC cells [80], and expression of ATF4 in fibroblasts was suggested to promote disease progression and resistance to chemotherapy in PDAC [61]. Other studies reported regulation of stress responses by *BRCA1. BRCA1* was shown to actively regulate reactive-oxygen-species (ROS) in response to oxidative stress [81], and to regulate the unfolded protein response (UPR)/ER stress response by regulating glucose-regulated protein (GRP)78, CHOP and GRP94 [82]. Furthermore, *BRCA1* induction led to downregulation of *HSF1* [83]. Our results indicate that while only *HSF1* was significantly higher in *BRCA*-mut tumors compared to *BRCA*-WT tumors, a broader stress network is activated in the *BRCA*-mut TME. This may reflect a mechanism by which DNA repair deficiencies in the cancer cells impose unique stress conditions on the TME that reshape the stress-response network in the stroma.

Several recent single-cell studies performed by us and others on human tumors and mouse models identified diverse CAF subtypes in breast, pancreatic, ovarian and prostate cancers [8, 10, 28, 84-86]. To which extent these subtypes are cancer-type specific or represent Pan-cancer markers is a burning question in our field. Even more pressing, perhaps, is the question of whether the mutation dependencies identified in our study are PDAC-specific or Pan-*BRCA*, or perhaps even represent a general characteristic of homologous recombination deficiency (HRD) cancers. These questions bear important implications on future therapeutic strategies. Defining common and segregating design principles of CAFs between tissues and organs sharing similar *BRCA*/HRD mutations will be an important step towards advancing therapy directed at these poor-prognosis cancers.

## Materials and methods

### Ethics statement

All clinical samples and data were collected following approval by Memorial Sloan Kettering Cancer Center (MSKCC; IRB, protocols #06-107, #15-015 and 13-217), Shaare Zedek Medical Center (IRB protocol #101/13; Ministry of Health no. 920130134), Sheba Medical Center at Tel-Hashomer (IRB protocol #0967-14-SMC), and the Weizmann Institute of Science (IRB, protocols # 186-1) Institutional Review Boards. All animal studies were conducted in accordance with the regulations formulated by the Institutional Animal Care and Use Committee (IACUC; protocol #01130121-2).

### Human patient samples

Tumor samples from surgically resected primary pancreas ductal adenocarcinomas were from patients treated at Memorial Sloan Kettering Cancer Center (MSKCC), at Shaare Zedek Medical Center, and at Sheba Medical Center; consent to study the tissue was obtained via MSK IRB protocols #06-107 and 13-217 for the main cohort (Cohort 1; Supplementary Table 1), and #15-015 for the exosome analysis (Cohort 2; Supplementary Table 6). Cohort 1 included a total of 27 *BRCA*-WT PDAC patients and 15 *BRCA*-mut PDAC patients (Supplementary Table 1). Cohort 2 included fresh samples from 26 patients from which tumor tissues and/or normal adjacent controls were collected (Supplementary Table 6). Of the 15 *BRCA*-mut patients, 4 are *BRCA1*-mut carriers and 11 are *BRCA2* carriers, which is consistent with the reported prevalence of *BRCA1* and *BRCA2* mutations in PDAC [87]. FFPE whole tumor sections and deeply annotated demographic, clinical, pathologic and genomic (MSK-IMPACT™) data were collected for all MSKCC patients in the study. In addition, fresh-frozen tumor tissue was collected for a subset of 12 patients.

### Mice

*Hsf1* null mice and their WT littermates (BALB/c × 129SvEV, by Ivor J. Benjamin [88]) were maintained under specific-pathogen-free conditions at the Weizmann Institute’s animal facility. Mice were sacrificed by CO_2_ for pancreata harvesting.

### Cell lines and primary cell cultures

**Mouse immortalized PSCs and KPC organoids and cell lines** were provided by David Tuveson’s laboratory [12]. Mouse PSCs were cultured in Dulbecco’s modified Eagle’s medium (DMEM; Biological industries, 01-052-1A) supplemented with 10% fetal bovine serum (FBS) and pen/strep.

**Primary PSC isolation:** pancreata were collected postmortem from *Hsf1* null mice or WT littermates into HBSS (Sigma-Aldrich, H6648), then minced into Roswell Park Memorial Institute 1640 (RPMI) (Biological industries, 01-100-1A), supplemented with 0.5mg/mL Collagenase D (Merck, 11088866001), 0.1mg/mL Deoxyribonuclease I (Worthington, LS002007) and 1mM HEPES (Biological Industries, 03-025-1B). Pancreata were incubated in 37 °C for 40 min with mechanical disruption every 5 min. Cells were then filtered with 100μm filters, centrifuged and isolated by Histodenz gradient (Sigma-Aldrich, D2158) dissolved in HBSS. Cells were resuspended in HBSS with 0.3% BSA and 43.75% Histodenz, HBSS with 0.3% BSA was layered on top of the cell suspension, and centrifuged for 20 min at 1,400 RCF. The cell band above the interface between the Histodenz and HBSS was harvested, washed in PBS, and plated in DMEM with 10% FBS and Pen/Strep. One week after seeding, immune and epithelial cells were depleted by anti-EpCAM (Miltenyi, 130-105-958) and anti-CD45 (Miltenyi, 130-052-301) magnetic beads, and transferred to LS columns (Miltenyi, 130-042-401). For gene expression measurements, PSCs were then cultured for 3 days and mRNA was isolated.

**Organoid lines derived from primary pancreatic KPC tumors** were provided by the laboratory of David Tuveson, and previously described [12]. KPC organoids were cultured in Corning® Matrigel® Growth Factor Reduced (GFR) Basement Membrane Matrix, Phenol Red-free, LDEV-free, (Corning, 365231) with complete organoid media [89]. Silencing of *Brca2* in KPC organoid lines was obtained by a lentiviral infection, using mouse *Brca2* shRNA in pLKO.1 vector (Horizon, RMM4534). Following viral infection, the organoids were selected for expression of shBRCA2 using puromycin. Conditioned media was collected following 3-4 days of culture with 5% FBS DMEM.

### PSC 3D cultures

For 3D culture, 2×10^4^ cells were seeded in Matrigel® GFR in DMEM supplemented with 10% FBS, L-glutamine and Pen/Strep for 4 days. Medium was changed to KPC conditioned medium or to their own conditioned media as control for 4 additional days and cells were harvested for RT-PCR. For HSF1 inhibition, 3 nM CMLD011866 (aglaroxin C) [67, 68] was added every 2 days.

### Immunohistochemistry of human tissues

4-µm FFPE sections from PDAC tumors were deparaffinized and treated with 1% H_2_O_2_. Antigen retrieval was performed and the antibodies listed in Supplementary Table 7 were used. Visualization was achieved with 3,3’-diaminobenzidine as a chromogen (Vector Labs, SK4100). Counterstaining was performed with Mayer hematoxylin (Sigma Aldrich, MHS16). Images were taken with a Nikon Eclipse Ci microscope (Figure 1A) or scanned by the Pannoramic SCAN II scanner, with 20×/0.8 objective (3DHISTECH, Budapest, Hungary) (Figure 2B-C).

### Multiplexed Immunofluorescent (MxIF) staining and imaging of human tissues

#### MxIF staining

FFPE sections from 22 *BRCA*-WT and 14 *BRCA*-mut PDAC patients were deparaffinized, and fixed. Antigen retrieval was performed. Slides were then blocked and used in a multiplexed manner with the OPAL reagents (AKOYA BIOSCIENCES). Briefly, following primary antibody incubation, slides were washed, incubated with secondary antibodies conjugated to HRP, washed again and incubated with OPAL reagents. Slides were then washed and antigen retrieval was performed. Then, slides were washed with PBS and stained with the next primary antibody or with DAPI at the end of the cycle. Finally, slides were mounted using Immu-mount (#9990402, Thermo Scientific).

#### MxIF imaging

FFPE samples were imaged with a LeicaSP8 confocal laser-scanning microscope (Leica Microsystems, Mannheim, Germany), equipped with a pulsed white-light and 405 nm lasers using a HC PL APO ×40/1.3 oil-immersion objective and HyD SP GaAsP detectors.

#### MxIF analysis

Images were analyzed using Fiji image processing platform [90]. For all panels of MxIF staining 3-5 images were obtained per patient. For each image, five slices were Z projected (max intensity) and linear spectral unmixing was performed. To assess the CAF composition (Figure 1) each channel was thresholded to create a mask of its area. The area of each CAF marker within CD45^-^ CK^-^ area was calculated by pixel-based analysis and divided by the total region of interest (ROI), as defined by CD45^-^ CK^-^ area. For the assessment of CAF subtype ratios, values were logged and averaged per patient. To study the discrete expression of the different CAF markers (Sup. Figure 1A-C) we performed an object-based analysis using the QuPath software [91]. CD45^+^ and CK^+^ cells were excluded. Then, the number of positive cells for each marker was calculated. The ratio of positive class cells for each marker (αSMA, CLU, HLA-DR) was defined as N/A if there were less than 10 positive cells of that marker in that image. If there were more than 10 positive cells within the image, all 1^st^ and 2^nd^ order overlap ratios (relative to the chosen marker) were calculated. All images per patient were averaged. To analyze TF activation (Figure 3) we used the Fiji image processing platform. First, we defined ROIs to exclude all cancer cells. Then, we detected nuclei of all stromal cells using the DAPI channel. Then, the mean intensity of each TF in all stromal cell nuclei per image was calculated. In case less than 10 cells were evident per channel, this channel was excluded per that image. For each patient, the average intensity of all images was calculated. To analyze the expression of MUC5B and SERPINA1 within CLU^+^ and αSMA^+^ CAFs we used Fiji image processing platform. First, we excluded all CK^+^ area. Then, the area of each of the CAF markers and secreted proteins was thresholded to create a mask of its area. Then, the area of CLU or αSMA out of MUC5B or SERPINA1 was calculated.

### Single cell validation

Human PDAC single-cell dataset (accession number: CRA001160) [14], was analyzed using the Seurat (V4.0) R toolkit [48, 92]. We analyzed all cells that were defined as ‘Fibroblast_cell’ or ‘Stellate_cell’ in the original dataset. Non-tumor samples and two samples with less than 50 Fibroblast or Stellate cells were filtered out. All other functions were run with default parameters. Harmony integration [93] with default parameter was used to minimize the patient batch effect and shared nearest neighbor (SNN) modularity optimization-based clustering was then used with a resolution parameter of 0.14 [94]. Two clusters were excluded from further analysis, one had less than 10 cells and the other is a cluster that expresses the pericyte marker, MCAM. Then we utilized the Seurat pipeline to analyze the remaining cells/patients. Pathway analysis was performed using Metascape [49]. UMAP images displaying gene expression were plotted using a minimum cutoff of the 10’th quantile.

### Pathway enrichment analysis

Pathway enrichment analysis was performed using Metascape [49] to analyze the DE genes in the RNA-seq results, as well as the different clusters of the single-cell data.

### Pixel classification of H&E stained slides from PDAC patient samples

H&E slides were scanned by the Pannoramic SCAN II scanner, with 20×/0.8 objective (3DHISTECH, Budapest, Hungary). Quantification of CAF-rich, cancer-rich, and immune-rich regions within the tumor area of each section was done by QuPath (version 0.2.3) [91] using pixel classification. The same classification parameters were used for all images.

### Laser capture microdissection of human PDAC samples

Fresh frozen blocks of *BRCA*-WT and *BRCA*-mut PDAC tumors were obtained from MSKCC (Supplementary Table 1). 7mm sections were sliced in a cryostat and placed on PEN Membrane Glass Slides (Thermo Fisher Scientific, LCM0522). Then, sections were stained using the Histogene™ LCM Frozen Section Staining Kit (Thermo Fisher Scientific, KIT0401) and stromal regions were dissected. Immune islands, cancer cells and blood vessels were excluded from microdissection. Slides were left to dry for 5 minutes at RT followed by microdissection using the Arcturus (XT) laser microdissection instrument (Thermo Fisher Scientific, #010013097). Infrared capture was used to minimize RNA damage. CapSure Macro LCM caps (Thermo Fisher Scientific, #LCM0211) were used to capture microdissected tissue. Microdissected tissue from each sample was incubated for 1 hr in 65 °C in the lysis buffer of the RNA extraction kit and frozen at -80 °C. RNA extraction was performed using the PicoPure™ RNA Isolation Kit (Thermo Fisher Scientific, KIT0204) according to manufacturer’s instructions.

### Library preparation and RNA-sequencing

Libraries were prepared using the SMARTer Stranded Total RNA-Seq v2-Pico Input Mammalian Kit (Takara Bio USA, #634415) according to the instructions of the manufacturer. Libraries were sequenced on Illumina NextSeq 500, at 25M reads per sample.

### Differential Expression Analysis

The DeSeq2 package [95] was used to identify differentially expressed genes between *BRCA*-mut and *BRCA*-WT samples, and the FDRtool [96] was used to compute local FDR values from the p-values calculated using DeSeq2. Genes with a fold change greater or equal to 1.5 and false discovery rate (FDR) of less than 0.05 were considered significantly differentially expressed between groups. Batch biases were corrected using Deseq2 package as RNA extraction and library preparation were performed in two batches.

### CIBERSORT

The abundance of immune cell populations in RNA-Seq samples was estimated using cell-type identification by estimating relative subsets of RNA transcripts (CIBERSORT) and LM22, a validated leukocyte gene signature matrix [50]. The quantile normalization was disabled as recommended by the CIBERSORT web interface (https://cibersort.stanford.edu/). Permutations were set to 500.

### Proteomic analysis of human exosomes

Fresh pancreatic cancer tissue and peritumoral non-involved pancreas tissues were cut into small pieces and cultured for 24 hours in serum-free RPMI, supplemented with penicillin (100 U/ml) and streptomycin (100 µg/ml). Conditioned media was processed for exosome isolation. Exosomes were purified by sequential ultracentrifugation as previously described [97, 98]. Briefly, cell contamination was removed from resected tissue culture supernatant by centrifugation at 500 x g for 10 min. To remove apoptotic bodies and large cell debris, the supernatants were then spun at 3,000 x g for 20 min, followed by centrifugation at 12,000 x g for 20 min to remove large microvesicles. Finally, exosomes were collected by ultracentrifugation twice at 100,000 x g for 70min. Five micrograms of exosomal protein were used for mass spectrometry analysis [98]. High resolution/high mass accuracy nano-LC-MS/MS data was processed using Proteome Discoverer 1.4.1.14/Mascot 2.5. Human data was queried against the UniProt’s Complete HUMAN proteome.

### Ingenuity

This tool generates multi-level regulatory networks by suggesting upstream regulators that may lead to the transcriptional changes evident in a dataset. To identify predicted upstream regulators of the BRCA-associated CAF transcriptional program we analyzed our differentially expressed gene dataset using the Causal Network tool in the Ingenuity Pathway Analysis (IPA) software [53].

### Expression analysis in the breast TCGA dataset

The TCGA public breast dataset was analyzed for expression of MUC5B in primary tumors from 745 *BRCA*-WT and 44 *BRCA*-mut patients.

### Real-time PCR

RNA was isolated using TRIzol reagent based on the TRI reagent user manual (Bio-Lab, 959758027100). Reverse transcription was done by High-Capacity cDNA reverse transcription kit (Cat 4368814, Thermo Fischer Scientific) according to the manufacturer instructions. Quantitative RT–PCR analysis was performed using Fast SYBR Green Master mix (Applied Biosystems, 4385610) or Taqman Fast Advanced Master Mix (Applied Biosystems, 4444556), and data was normalized to the house-keeping gene HPRT. Primer sequences for qPCR used in this study are provided in Supplementary Table 8.

### Aglaroxin C

((-)-Aglaroxin C (CMLD011866) was synthesized according to published protocols [67, 68] and used in a concentration of 3nM.

### Chromatin immunoprecipitation (ChIP) followed by qRT-PCR

Immortalized PSCs were treated with KPC conditioned medium. 24 hrs later, PSCs were harvested for CHIP assay as described in [99]. Anti-HSF1 Ab (Cell Signaling, 4356S) was used to immunoprecipitate HSF1, and normal rat IgG (Cell Signaling, 2729S) was used as control. qRT-PCR was performed to assess the binding of HSF1 to *Clu, Hsp90aa*, and *Hspa1a* with genomic primers designed to flank an HSE site homologues to that reported to bind HSF1 in human cells [56]).

### Conditioned medium for Chromatin IP

KPC cells were plated in a density of 15×10^4^/cm^2^ in DMEM supplemented with 5% fetal bovine serum (FBS) and L-glu and Pen/Strep. 24h later, the medium was replaced and cells were left to grow for an additional 48h. The medium was then collected and filtered through 0.22μm filters, and placed on top of PSC cultures.

### Statistical analysis

Statistical analysis and visualization were performed using R (Versions 3.6.0 and 4.0.0, R Foundation for Statistical Computing Vienna, Austria) and Prism 9.1.1 (Graphpad, USA). Statistical tests were performed as described in each Figure legend. Mann Whitney test was used to analyze data that is not normally distributed. Student’s t-test or ANOVA were used to analyze normally distributed data. Pearson’s correlation coefficient was used to assess the association between two continuous variables. All statistical tests were defined as significant if *p* value < 0.05 or FDR< 0.05 for multiple comparisons. “ns” in all Figure s marks p-values greater than 0.05. Values that were more than 2 standard-deviations of their group mean were defined as outliers and excluded.

## Data availability

RNA-sequencing data will be deposited in the dbGaP Authorized Access System. The remaining data are available within the Article, Supplementary Information or available from the authors upon reasonable request.

---

**Figure S1.**
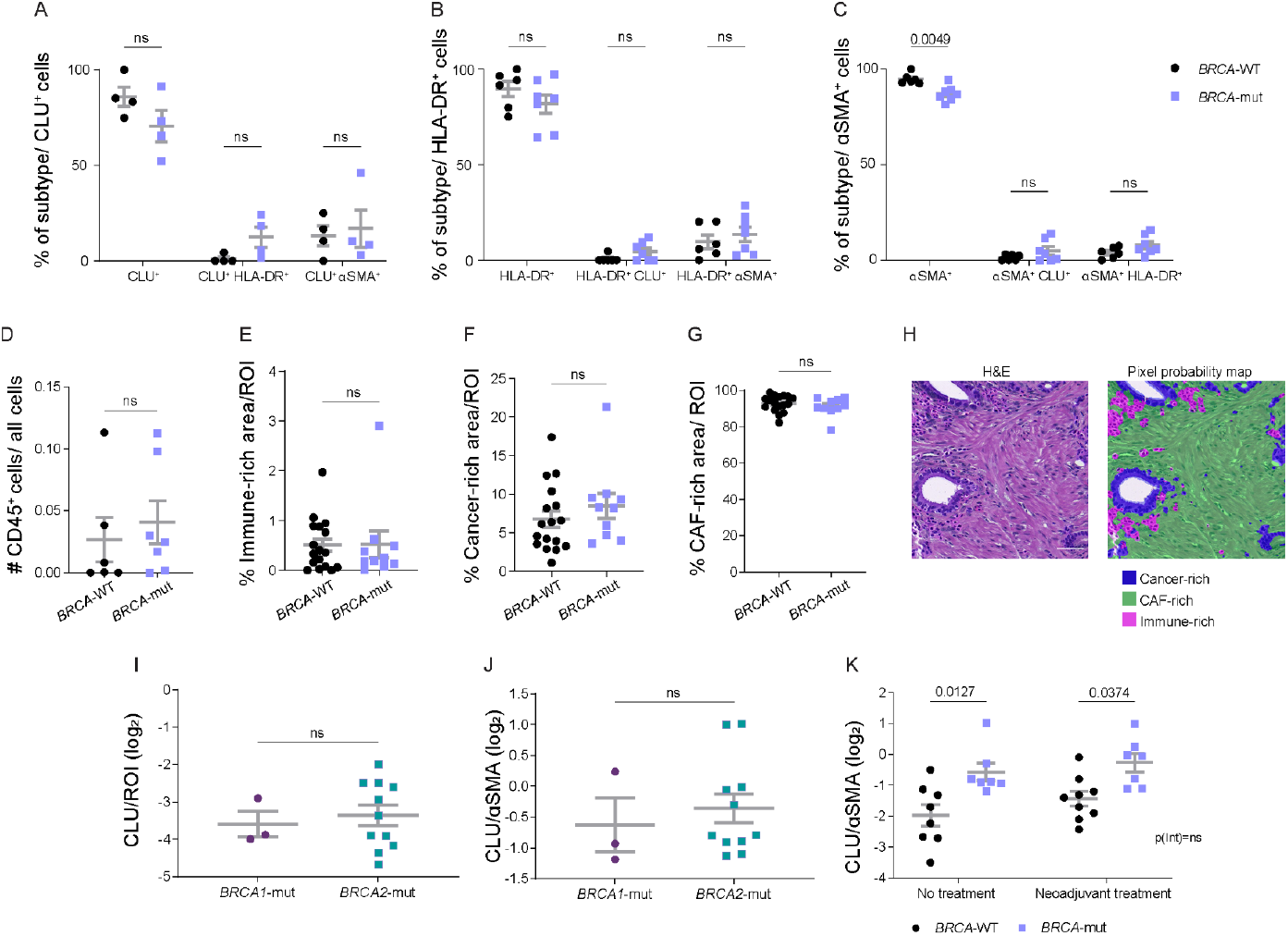
PDAC stroma is comprised of three distinct CAF subtypes. (A-C) MxIF was performed on FFPE tumor sections from 7 *BRCA*-mut and 6 *BRCA*-WT PDAC patients using antibodies for αSMA, CLU, HLA-DR, CK, and CD45. DAPI was used to stain nuclei. Images were analyzed with QuPath software using object-based analysis to detect different CAF subtypes and cell markers. The fraction of each subtype out of all the cells that expresses CLU (A), HLA-DR (B), and αSMA (C) is presented. (D) The relative abundance of CD45^+^ cells was calculated, averaged for each patient sample, and is presented as mean ± SEM across patients. (E-H) 17 *BRCA*-WT and 10 *BRCA*-mut H&E stained FFPE slides were scanned and analyzed employing pixel classification using QuPath software. The percent of immune-rich (E), cancer-rich (F), and CAF-rich (G) areas out of an ROI of the tumor area (excluding normal adjacent tissue) was calculated. Representative H&E image before (left) and after (right) pixel classification are shown in (H). Blue marks cancer-rich areas, green marks CAF-rich areas, and pink marks immune-rich. Scale bar - 50 µm. (I-J) FFPE tumor sections from 17 *BRCA*-WT and 14 *BRCA*-mut carriers (n=11 for *BRCA2* and n=3 for *BRCA1*) PDAC patients were stained by MxIF for CLU, αSMA, and cytokeratin (CK). DAPI was used to stain nuclei. Images were analyzed using ImageJ software, CK^-^ regions were defined as regions of interest (ROIs) and the area stained by each CAF marker was calculated, divided by the ROI and averaged for each patient sample. Quantification of the CLU^+^ area (I) and the ratio between CLU^+^ and αSMA^+^ CAFs (J) is presented for tumor samples from *BRCA1 vs. BRCA2* mutation carriers. Quantification of the ratio between CLU^+^ and αSMA^+^ CAFs for tumor samples from neoadjuvant-treated *vs*. non-treated patients is presented (K). Data are presented as mean ± SEM. ns marks p-values greater than 0.05.

**Figure S2.**
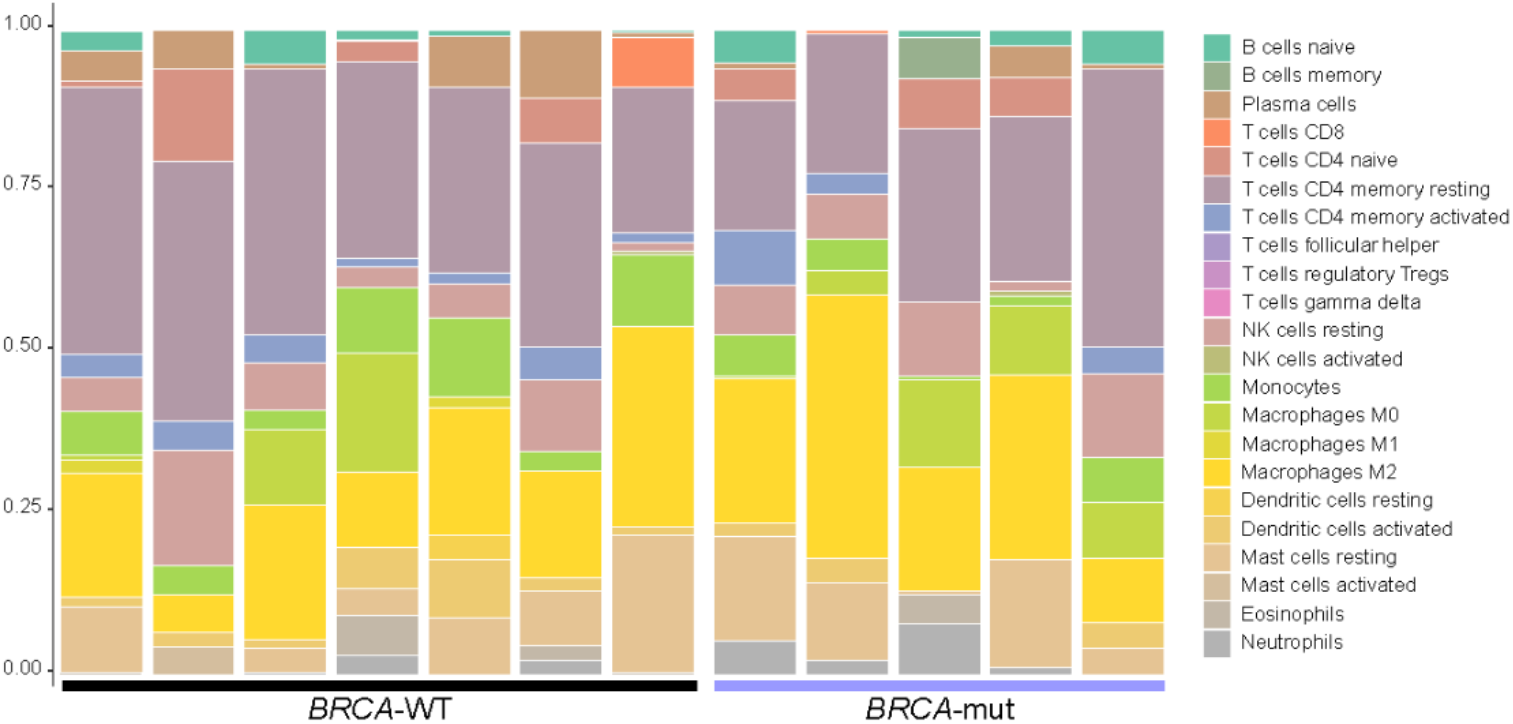
CAF-rich regions from *BRCA*-WT and *BRCA*-mut PDAC show similar immune compositions. RNA-seq data from laser capture microdissected CAF-rich regions of 7 *BRCA*-WT and 5 *BRCA*-mut PDAC patients was analyzed using CIBERSORT to estimate immune cell composition of *BRCA*-mut and *BRCA*-WT CAF-rich regions. The predicted fraction of each immune cell population is presented.

**Figure S3.**
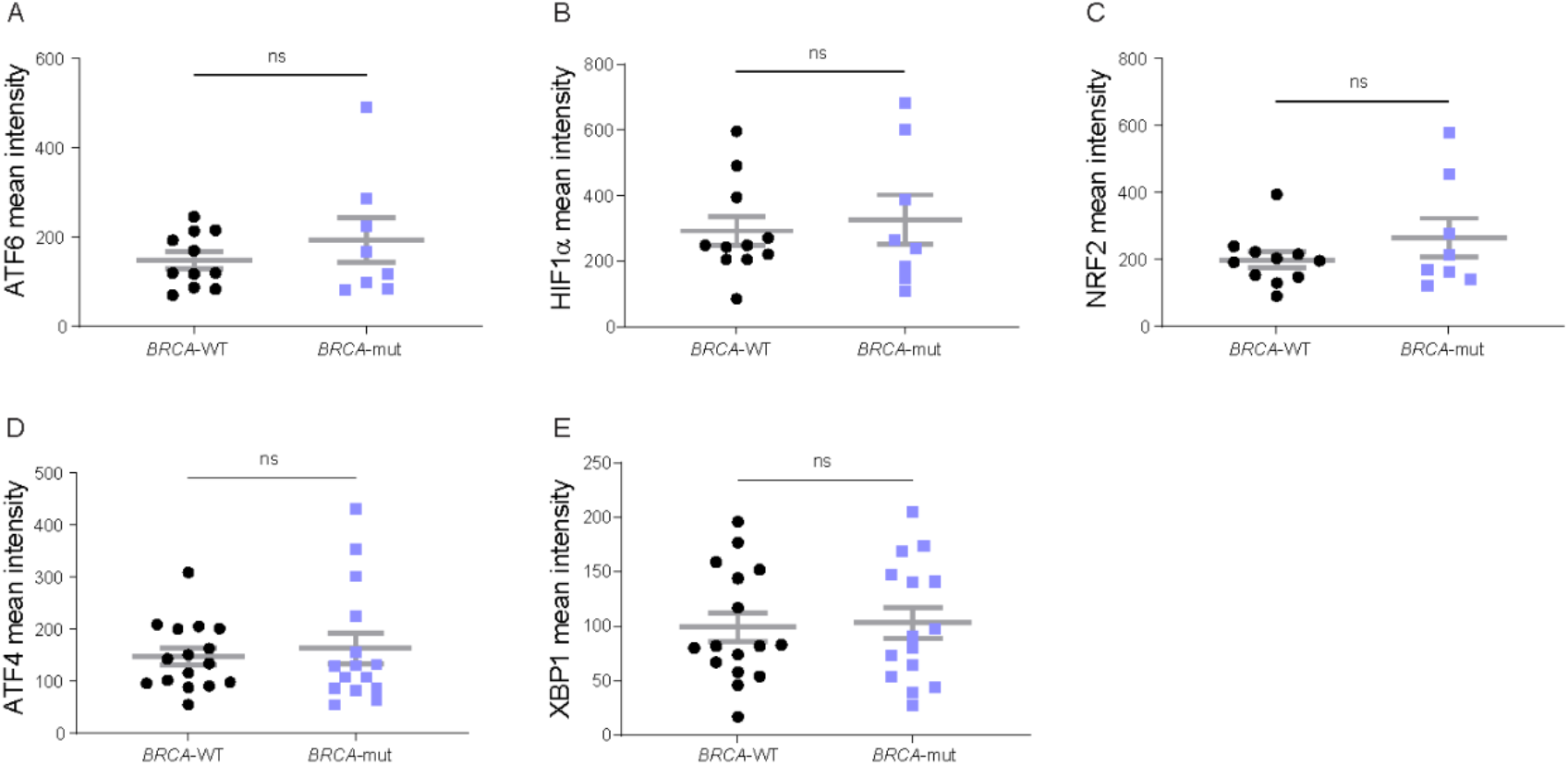
CAFs from *BRCA*-WT and *BRCA*-mut PDAC exhibit similar activation levels of the responses to ER stress, integrated stress, hypoxia, and oxidative stress. FFPE tumor sections from *BRCA*-mut and *BRCA*-WT PDAC patients were stained by MxIF using antibodies for the indicated proteins. 3-5 images per patient were analyzed using ImageJ software. (A-E) Quantification of the nuclear staining (mean intensity) of ATF6 (A), HIF1a (B), NRF2 (C), ATF4 (D), and XBP1 (E) within all stromal cells in *BRCA*-mut (n=8 for A-C; n=15 for D-E) and *BRCA*-WT samples (n=11 for A-C; n=16 for D-E). Data are presented as mean across patients ± SEM. ns marks p-values greater than 0.05.

**Figure S4.**
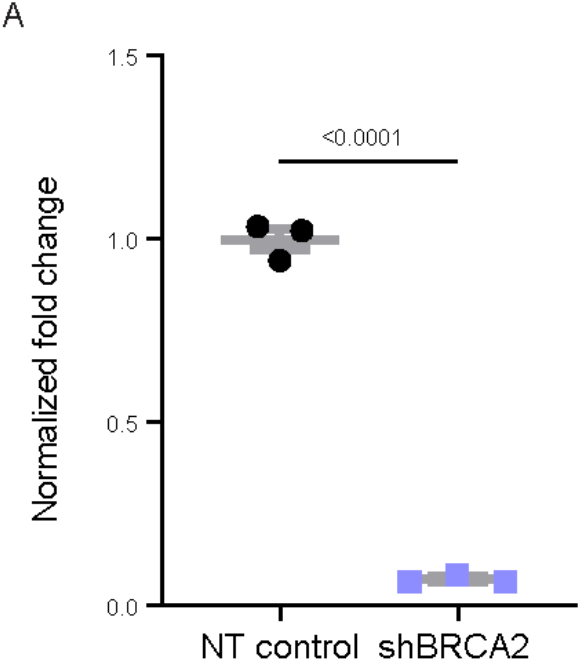
Validation of *Brca*2 knockdown by shRNA in KPC organoids. KPC organoids were transfected with sh*Brca*2. *Brca*2 levels were measured by qRT-PCR compared to non-targeting shControl. Data are presented as mean ± SEM. n=3 technical repeats per group.

